# Landmark-Based Navigation Instructions Improve Incidental Spatial Knowledge Acquisition in Real-World Environments

**DOI:** 10.1101/789529

**Authors:** Anna Wunderlich, Klaus Gramann

## Abstract

The repeated use of navigation assistance systems leads to decreased processing of the environment. Previous studies demonstrated that auditory augmentation of landmarks in navigation instructions can improve incidental spatial knowledge acquisition when driving a single route through an unfamiliar virtual environment. Based on these results, three experiments were conducted to investigate the generalizability and ecological validity of incidental spatial knowledge acquisition by landmark augmentation in navigation instructions.

In the first experiment, spatial knowledge acquisition was tested after watching an interactive video showing the navigation of a real-world urban route. A second experiment investigated incidental spatial knowledge acquisition during assisted navigation when participants walked through the same real-world, urban environment. The third experiment tested the acquired spatial knowledge two weeks after an assisted pedestrian navigation phase through the real-world.

All experiments demonstrated better performance in a cued-recall task for participants navigating with landmark-based navigation instructions as compared to standard instructions. Different levels of information provided with landmark-based instructions impacted landmark recognition dependent on the delay between navigation and test. The results replicated an improved landmark and route knowledge when using landmark-based navigation instructions emphasizing that auditory landmark augmentation enhances incidental spatial knowledge acquisition, and that this enhancement can be generalized to real-life settings.

This research is paving the way for navigation assistants that, instead of impairing spatial knowledge acquisition, incidentally foster the acquisition of landmark and route knowledge during every-day navigation.

## Introduction

Navigation aids have become common everyday tools (Axon et al., 2012; Kalin & Frith, 2016; Kitchin & Dodge, 2007). They provide visual as well as auditory guidance as a support during wayfinding through known and unknown environments (Allen, 1999). With the use of navigation aids, wayfinding evolved from analog 2D map-based tasks into digitally assisted instruction following tasks.

Several studies investigated the user’s interaction with navigation aids in order to describe and understand the underlying cognitive processes. One finding showed that the use of automated assistance systems was associated with divided attention between the movement related task and the assisted navigation (Fenech et al., 2010; Gardony et al., 2013; Gardony et al., 2015). This resource allocation conflict increases the reliance on the navigation assistant in order to reduce attentional demands (Baus et al., 2001; Klippel et al., 2010; Parush et al., 2007) leading to an automation bias (Lin et al., 2017). The users tend to hand over the decision-making to the automated system (Bakdash et al., 2008; Fenech et al., 2010; Parush et al., 2007). They follow the system instructions without checking other available information (Mosier et al., 1996; Parasuraman, 2000) and this over-reliance leads to a decrease in the processing of the surrounding environment (Fenech et al., 2010; Hirtle & Raubal, 2013; Leshed et al., 2008).

As a consequence, when the navigation aid is no longer reliably functioning (e.g., because of system errors or GPS signal loss), people are more likely to get lost because of their inability to extract navigation relevant information from the environment and to orient using their own sense of direction. However, even when the system works properly, the risk of getting lost remains based on inadequate application of or over-reliance on the navigation assistance system. This was revealed in in a corpus of 158 so called “Death by GPS” incidents published in the English news between 2010 and 2017 (Lin et al., 2017). Only those reports were chosen that contained incidents caused by an improper use of navigation assistance systems reporting an unusual high number of single-car crashes (32%) and that most of the “Death by GPS” incidents happened in an unknown environment (78%). These data point to a relative increase of rare accidents involving no other road users demonstrating that the use of automated assisting technologies can negatively impact the primary driving task.

Thus, a first goal of an improved navigation assistance system should be to maintain the ability to extract navigation relevant information from the environment without endangering the users’ safety. One solution is the inclusion of environmental information about salient objects, so called landmarks (Evans et al., 1982), in the navigation instructions (Goodman et al., 2005; Li et al., 2014; May et al., 2001). It was shown that landmark knowledge can be incidentally acquired during navigation to a similar level as intentional learning (Chrastil & Warren, 2012; Van Asselen et al., 2006). Compared to visual augmentation methods, acoustic navigation instructions have the advantage to not interfere with visual attention necessary to observe the ongoing traffic (May & Ross, 2006). Ross et al. (2004) investigated landmark-based auditory navigation instructions regarding their usability for pedestrians. The authors demonstrated the effectiveness of landmark-based auditory instructions leading to fewer navigation errors and increased navigator’s confidence during navigation, as compared to a control group. While the study by Ross and colleagues demonstrated improved navigation performance, the authors did not test for the impact of auditory landmark information on spatial knowledge acquisition.

To test whether auditory landmark-based navigation instructions lead to increased processing of environmental information beyond improving save navigation, Gramann et al. (2017) augmented landmarks at intersections in a virtual driving task. Landmark augmentation was implemented by naming landmarks and providing additional information about the landmark at intersections with route direction changes. This resulted in incidental acquisition of navigation relevant information about the environment (Gramann et al., 2017). This improved landmark and route knowledge acquisition with landmark-based navigation instructions was observable even when tested three weeks after a single exposure to an unfamiliar environment (Wunderlich et al., 2020). It was further found to be associated with changes in brain activity likely reflecting increased information recollection during cued-recall of landmark- and route-knowledge about the navigated environment in general (Wunderlich & Gramann, 2018).

The results of both virtual driving studies, Gramann and colleagues (2017) as well as Wunderlich and Gramann (2018), showed a significantly improved recognition performance for landmarks at intersections with route direction changes when navigators received landmark-based instructions as compared to standard navigation instructions known from commercial systems. These results support the assumption that the inclusion of landmark information in navigation instructions was associated with directing the users’ attention towards environmental features. Landmarks (i.e., salient and lasting aspects of the surrounding environment like buildings) are essential elements of spatial representations and necessary for conceiving spatial relations (Ekstrom & Isham, 2017; Siegel & White, 1975). Augmenting landmarks in auditory navigation instructions might thus be a promising way to foster processing of the environment leading to incidental spatial knowledge acquisition during the use of navigation assistance systems. Importantly, the enhanced processing of the environment when using landmark-based instructions did not impact the subjective mental load or the driving behavior, thus, securing safety of the primary driving task (Gramann et al., 2017; Wunderlich & Gramann, 2018).

The reported studies by Gramann and colleagues (2017) as well as Wunderlich and Gramann (2018) used two versions of landmark-based navigation instructions that included the name of a landmark at the intersection and additional landmark information. One instruction condition added personally relevant information to landmarks (for example: “Turn right at the bookstore. There, you can buy books of J.R.R. Tolkien” in case J.R.R. Tolkien was the favorite author of the tested participant). Personal interests in different categories were acquired prior to the experiment and navigation instructions were individualized for each participant accordingly. A second instruction condition provided redundant information of comparable length as the personal-reference condition and was identical for all participants in this group (e.g., “Turn right at the bookstore. There, you can buy books.”) To answer the question whether the information with personal interest or the more detailed information was the cause for the potential differences in acquired spatial knowledge, the here presented studies used a *long navigation instruction condition* that named landmarks and provided additional semantic information about the landmark (Wunderlich et al., 2020). This instruction contained more information than the previous redundant description but had no relation to personal interests (e.g., “Turn right at the bookstore. There, public readings take place every week.”). Because previous studies revealed improved spatial knowledge acquisition for any landmark-based navigation instructions, we also introduced a *short navigation instruction condition* to test whether simply naming a landmark without additional information might already foster spatial knowledge acquisition (“Turn right at the bookstore.”). This condition was similar to the landmark-based navigation instructions in the usability study by Ross et al. (2004). In summary, in three experiments, both short and/or long landmark-based navigation instructions were compared with a control group that received standard auditory navigation instructions as known from commercial navigation aids referring to the next intersection (e.g., “At the next intersection turn right.”).

Besides investigating the impact of landmark information in navigation instructions on the learning of the environment, it remained an open question whether the improved incidental spatial knowledge acquisition that was observed for driving simulations would generalize to more ecologically valid scenarios. The previous experiments took place in a driving simulator setup (Gramann et al., 2017; Wunderlich & Gramann, 2018) that lacked naturalistic self-motion cues, other traffic participants, and natural visual information. To investigate whether incidental spatial knowledge acquisition with landmark-based navigation instructions can be observed also for more realistic scenarios, we conducted three experiments. In experiment 1, we transferred the navigation paradigm to a pedestrian context using real-world visuals. An interactive video was created showing a pedestrian’s first-person perspective while navigating through Berlin, Germany. The video was a realistic representation of city navigation while allowing control of stimulus material for all participants. Experiment 2 and 3 investigated incidental spatial knowledge acquisition during real-world assisted pedestrian navigation through the city of Berlin. Experiment 2 investigated the identical route and stimuli that were used in Experiment 1 (Wunderlich & Gramann, 2020), whereas Experiment 3 extended the route used in experiments 1 and 2 to increase the number of intersections. In addition, the last experiment introduced a break of two weeks between the navigation phase and the subsequent spatial tests similar to Wunderlich and Gramann (2018).

We expected to observe the same positive impact of landmark-based navigation instructions on processing of the environment and related incidental spatial knowledge acquisition in both, the controlled video scenario as well as the real-world setting. Furthermore, we expected that participants receiving long instructions outperform those receiving short navigation instructions and hypothesized that landmark-based navigation instructions lead to a generally increased processing of the environment (also during straight segments) rather than only for auditory augmented landmarks at intersections (*halo effect*).

## Experiment 1

The aim of Experiment 1 was to test landmark-based navigation instructions in a more realistic environment providing real-world visuals. To this end, an interactive video was created showing a pedestrian’s first-person perspective during real-world navigation through the city of Berlin as an example for an urban environment.

### Method

#### Participants

The data of forty-three participants, assigned to three different navigation instruction groups, was evaluated in this experiment. All participants had normal or corrected to normal vision and gave written informed consent prior to the study. The study was approved by the local ethics committee. The sample included 22 women and gender was balanced across navigation instruction conditions (14 participants in standard, 7 females; 14 in short, 7 females; and 15 in the long instruction condition, 8 females). The age ranged from 19 to 34 years (*M* = 26.2 years, *SD* = 2.75 years). Participants were recruited through an existing database or personal contact and were reimbursed monetarily or received course credit. To assure that participants were unfamiliar with the route, we used an online questionnaire prior to the experiment. Participants were asked to rank up to five metro stations of Berlin according to their frequency of personal use. If one of the stations was close to the route, the participant was excluded from the experiment. After the navigation task, the participants were asked to rate their familiarity with the navigated route on a scale ranging from 0% (completely unknown) to 100% (completely known) and if their response exceeded 50% familiarity they were excluded from analyses before they proceeded with the spatial tasks. The reported sample had an averaged familiarity score of *M* = 9.88% (*SD* = 11.1%, standard: *M* = 9.43%, *SD* = 11.3%, short: *M* = 10.0%, *SD* = 13.6%, long: *M* = 10.2%, *SD* = 9.06%, *F* _(2,40)_ = 0.02, *p* = .983).

#### Procedure

Following previous experimental protocols (Gramann et al., 2017; Wunderlich & Gramann, 2018), the experiments consisted of two parts. The first part was an assisted navigation phase in which participants followed auditory navigation instructions to navigate along a predefined route. In the second part, participants had to solve different tasks testing their spatial knowledge about the navigated environment. Participants were not informed prior to the second part that they would be tested on the navigated environment. The entire experiment took place in a controlled laboratory environment (Figure 1a) and lasted 2 hours and 40 minutes. Participants were seated in front of a display and interacted with the video scenario by using the keyboard in front of them. Participants were equipped with electroencephalography (EEG; BrainAmps, Brain Products, Gilching, Germany) and eye movements were recorded with a desktop-based eye-tracker (SMI RED 5, SensoMotoric Instruments, Teltow, Germany).

**Figure 1.**
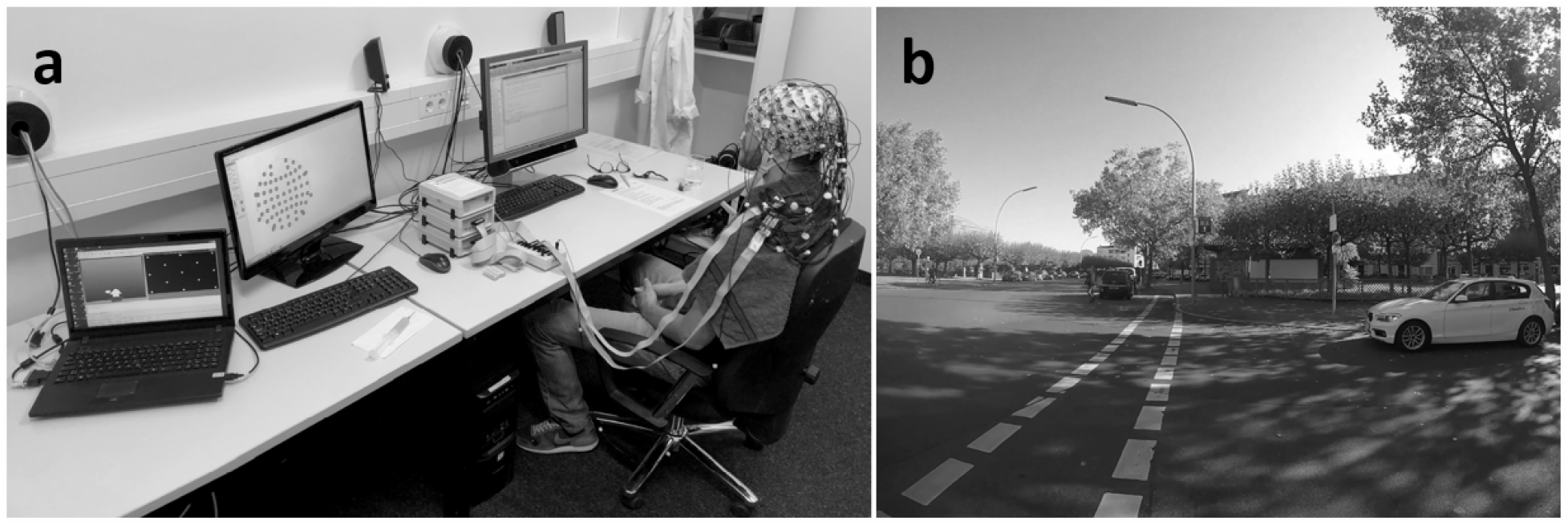
a) Interactive video setup of Experiment 1. The screen in front of the participant displayed the video and was equipped with the eye-tracker, a second screen was used to check the raw EEG-data, and the laptop for eye-tracking data collection. b) Video snapshot of a landmark at an intersection (“public restroom”) as seen during video-based navigation as well as used in the cued-recall task (pictures were presented in color).

#### Navigation Phase

Participants watched a video showing real world recordings complemented by auditory navigation instructions. The task was interactive as the video stopped at intersections and participants had to respond according to the navigation instructions. The video did not commence without a response. Due to the paradigm shift from simulated driving to walking through an urban environment, navigation instructions were provided prior to all intersections and landmarks at intersections were referenced in the landmark-based navigation instructions. In the previous studies, landmarks at intersections with a change in route direction were labelled navigation relevant while landmarks at intersections without direction change and no instruction were labelled navigation irrelevant. For experiment 1 and all following experiments, this definition was adapted to the pedestrian context that provided more time between intersections as compared to simulated driving. Participants received auditory navigation instructions prior to each intersection. Thus, landmarks located at *intersections* were merged into one category irrespective of whether the route remained straight or turned left or right.

Landmarks located along *straight segments* between intersections were considered the second landmark category. Both types of landmarks at intersections and along the route can be perceived as navigation relevant. While landmarks at intersections can be associated with changes in route direction, it was shown for pedestrian navigation assistance that landmarks located along straight route segments allow for confirming correct route progress and improve user confidence during navigation (May et al., 2003; Rousell & Zipf, 2017). However, the present experiments provided landmark-based instructions only for landmarks at intersections. Landmarks in between intersections were not accompanied by navigation instructions.

The video content for the navigation task was recorded with a GoPro Hero 4 (GoPro. Inc., San Mateo, USA) from a pedestrians’ perspective (see Figure 1b). The camera person was continuously recording while walking a predefined route. During the video, various other road users were visible. Removing the real-world audio allowed for high realism of the visuals without considering other auditory stimuli than the auditory navigation instructions. The video was accelerated by 1.1 times the original speed to shorten the navigation task to approximately 35 minutes without distorting the navigation experience. The displayed route through Charlottenburg, Berlin in Germany, was 3.7 km long and passed twenty intersections containing 14 route direction changes. Auditory navigation instructions were presented via loudspeakers prior to each intersection. When arriving at an intersection, the video stopped and a participants’ response according to the instructed direction was required. Participants indicated the navigation directions using the left (turn left), up (go straight), or right (turn right) arrow key. This interaction was implemented to ensure the participants’ attention to the instructions and the route. Irrespective of whether the response was correct or not, pressing one of the response keys started the video again. The stimulus material was identical for all participants and only the navigation instructions differed between the groups. Besides the standard navigation instructions, the short and long landmark-based navigation instructions were tested. After the video navigation phase, the subjectively experienced mental load was measured using the NASA-TLX (Hart, 2006; Hart & Staveland, 1988). In addition, participants answered questions about route familiarity and acceptance of the modified navigation instructions.

#### Test Phase

The test phase followed the navigation phase after a short break (as in Gramann et al., 2017). This second part consisted of sketch map drawing, the cued-recall task, the Perspective Taking/Spatial Orientation Task (PTSOT; Hegarty and Waller, 2004), and the circle task (triangular pointing task). Questionnaire data about demographic characteristics, individual navigation and learning habits, as well as spatial abilities were collected at the end of the experiment. The self-reported sense of direction was rated using 7 point Likert scale for agreement with the key item "My sense of direction is very good" (SOD, Hegarty et al., 2002; Kozlowski and Bryant, 1977).

Only the cued-recall task is reported in detail as it was the only task that was used in all three experiments included here as well as in the previous studies. This way, five studies could be directly compared and the results allowed for conclusions about the generalizability of the impact of landmark-based navigation instructions for different navigation contexts.

#### Cued-Recall Task

The cued-recall task combined a simple landmark recognition with a route direction response comparable to Huang et al. (2012). The task of the participant was to correctly recognize pictures taken from the navigated environment as being part of or not being part of the previously navigated route. In case of recognizing a landmark, participants were instructed to respond according to the respective route direction at this landmark location (straight, left, right). To capture the overall acquired landmark and route knowledge, we used three landmark types which were categorized according to their location: a) landmarks at *intersections* that were referenced in the modified navigation instructions, b) landmarks located at *straight segments* along the route, and c) *novel* landmarks that were part of the environment, but had not been encountered during the navigation phase (e.g., buildings in a parallel street). Thus, landmarks at intersections and landmarks along straight segments of the previously navigated route should be indicated as known landmarks by responding with the respective route direction while novel landmarks (independent of location) should be identified as unknown. Half of the novel landmarks were located at intersections while the other half was located alongside straight segments of a street that provided no turning opportunities. We analyzed response rates for known and novel landmarks at intersections as well as at straight segments of the route using signal detection theory to compute the sensitivity measure d’ (Atkinson, 1963; Marcum, 1960). This measure allowed to better control for possible response biases as discussed in Wunderlich and Gramann (2018) by considering the false alarm rate for novel landmarks of the respective landmark type. Comparing the sensitivity across navigation instruction conditions allowed for investigating differences in the acquired landmark knowledge. To investigate the incidentally acquired route knowledge, the performance in a cued-recall task was analyzed as a second dependent measure using the percentage of correct direction responses regarding the landmarks at intersections. This measure represents a combination of landmark and route knowledge of the navigated environment because correct responses required associative memory of the respective route direction. A higher percentage of correct responses represents more route knowledge that can be recollected based on the presented picture cues.

In the cued-recall task of experiment 1, twenty pictures of each landmark type (intersections, straight segments, novel) were randomized and presented one by one on a desktop screen. Pictures of intersections and straight segments were screenshot from the navigation video. Screenshots from additional video material which was taken the same day as the navigation video served as novel landmarks. All landmarks were presented from the pedestrians’ point of view including their immediate surroundings (see Figure 1b). Each landmark was displayed for 3 seconds and then replaced by four arrow keys indicating the response options. Participants were instructed to press the right/left arrow key, in case the route had turned right/left at the displayed landmark, respectively. In case the route had proceeded straight ahead, the required response was to press the up-arrow key. In case the landmark had not been encountered previously, participants were asked to press the down-arrow key. The location of landmarks relative to the intersections (left, right, before or after the intersection) did not provide any indication for the respective turning directions. Following each response, participants were asked how confident they were about their given answer on a six-point scale ranging from ‘1 - very sure’ to ‘6 - I did not know’ and a seventh option to state ‘I know that my last response was wrong’.

#### Statistical Analysis

Statistical analysis was performed using the statistics software SPSS (International Business Machines Corporation (IBM) Analytics, Armonk, USA).

To control for an impact of individual differences on the dependent variables, the percentage of correct responses to each landmark type and sensitivity values were correlated with the self-reported SOD and the averaged error in the PTSOT to test for a covariance of performance and subjective or objective spatial abilities, respectively. The same way, we also tested a potential influence of subjective mental load during the navigation task. In case of significant correlations with performance in the cued-recall test, the measures were used as covariate in the respective analysis.

The mixed-measure analysis of variance (ANOVA; ANCOVA in case of significant correlations with the additional measures) testing the incidentally acquired landmark knowledge used the between-subject factor navigation instruction condition (*standard, short, long*) and two levels for the factor landmark type. The first, *intersections*, counted all responses as a hit that correctly indicated that the landmark was encountered during navigation. This response represents the recognition sensitivity for intersection landmarks. The second, *straight segments*, represented the sensitivity to the landmarks along the route.

Incidentally acquired route knowledge was tested using an ANOVA (ANCOVA in case of significant correlations with the additional measures) for the percentage of correct responses to landmarks at intersections. The between-subject factor was navigation instruction (*standard, short, long*).

Post-hoc comparisons were computed comparing the instruction conditions and p-values were corrected for multiple comparisons using Bonferroni. As an indicator of effect size, partial eta squared was calculated.

### Results

#### Individual Measures

Individual measures of the PTSOT revealed a mean score of *M* = 26.7° (*SD* = 22.9°, standard: *M* = 23.2°, *SD* = 15.4°, short: *M* = 29.1°, *SD* = 25.1°, long: *M* = 27.5°, *SD* = 27.3°, *F* _(2,40)_ = 0.24, *p* = .787). The SOD rating was *M* = 3.88 (*SD* = 1.56, standard: *M* = 3.93, *SD* = 1.82, short: *M* = 3.43, *SD* = 1.60, long: *M* = 4.27, *SD* = 1.22, *F* _(2,40)_ = 1.05, *p* = .358) and the subjective mental load during assisted navigation rated on a scale from 1 (low) to 100 (high) *M* = 21.2 (*SD* = 18.0, standard: *M* = 21.4, *SD* = 24.3, short: *M* = 16.2, *SD* = 11.1, long: *M* = 25.7, *SD* = 16.1, *F* _(2,40)_ = 1.00, *p* = .375).

The Spearman correlation of the SOD rating with the dependent variables was significant for d’ of landmarks at intersections (*ρ* = .423, *p* = .005), other |*ρ*|’s < .193, *p*’s > .217). The averaged error of the PTSOT correlated negatively with d’ for both, landmarks at intersections (*ρ* = −.307, *p* = .046) and those at straight segments (*ρ* = −.312, *p* = .042), the correlation with percentage correct responses to landmarks at intersections was not significant (*ρ* = −.100, *p* = .523). No correlation of subjective mental load was significant (all *ρ*’s < .210, *p*’s > .177). The mean error of the PTSOT was included as a covariate in the 2×3 ANOVA testing the recognition sensitivity of the participants due to its significant correlation with both factor levels.

#### Landmark Knowledge

The sensitivity values can be seen as group-boxplots and individual measures in Figure 2. The results of the ANCOVA testing recognition sensitivity with the PTSOT error as covariate revealed a significant main effect of the covariate (*F*_(1,39)_ = 8.37, *p* = .006, *η*_p_^2^ = .177). The interaction of landmark type and navigation instruction condition did not reach significance (*F*_(2,39)_ = 2.51, *p* = .095, *η*_p_^2^ = .114). The main effect of navigation instruction condition and its interaction with the covariate PTSOT were also not significant (all *p*’s > .137).

**Figure 2.**
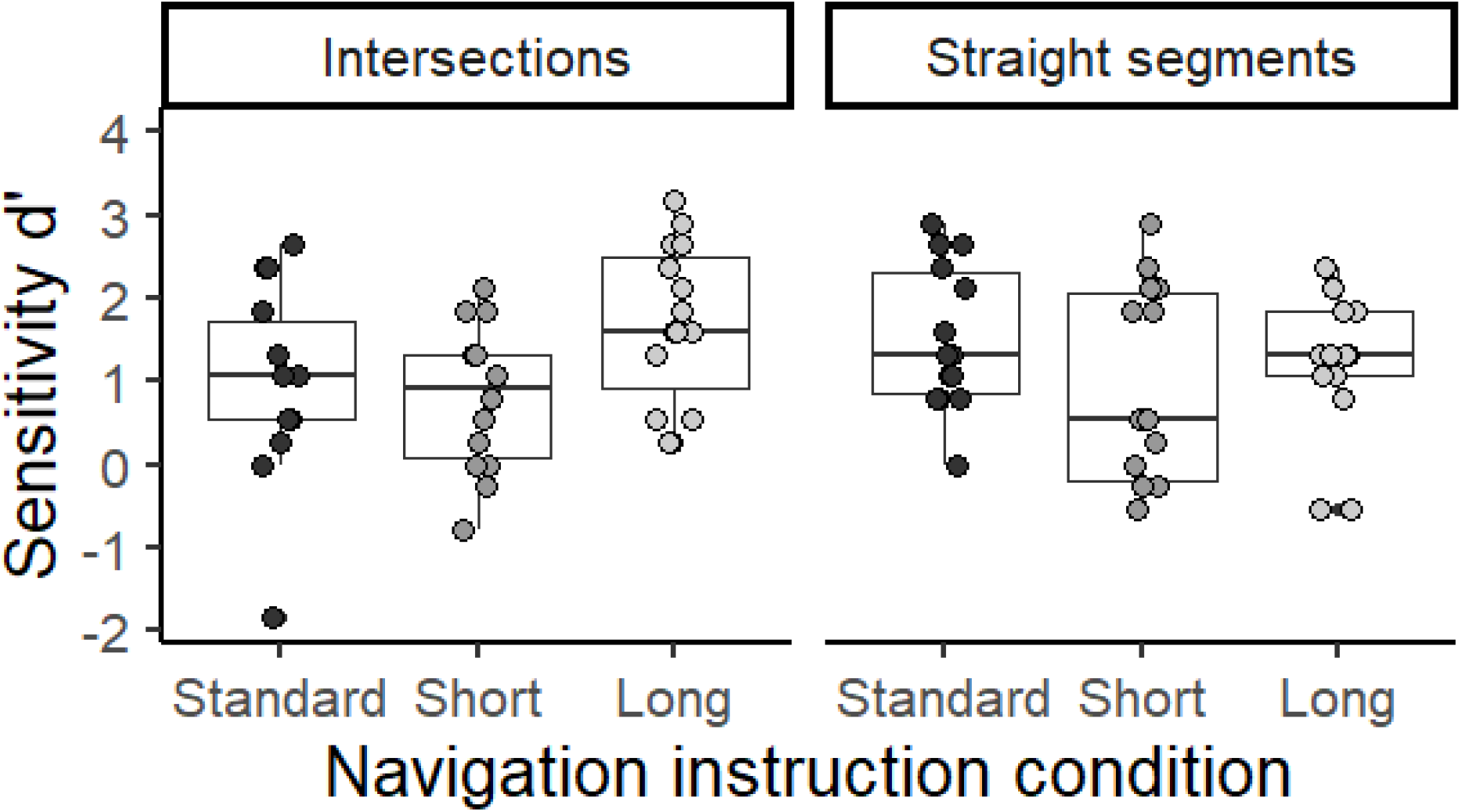
The sensitivity for correctly recognizing familiar landmarks was presented in box plots and single subject values of d’ for Experiment 1. The d’ represents the z-standardized rate of hits subtracted by the z-standardized rate of false alarms. Separated in columns, the sensitivity was plotted for landmarks at intersections and at straight segments. Small dots represent individual values above the whiskers.

**Figure 3.**
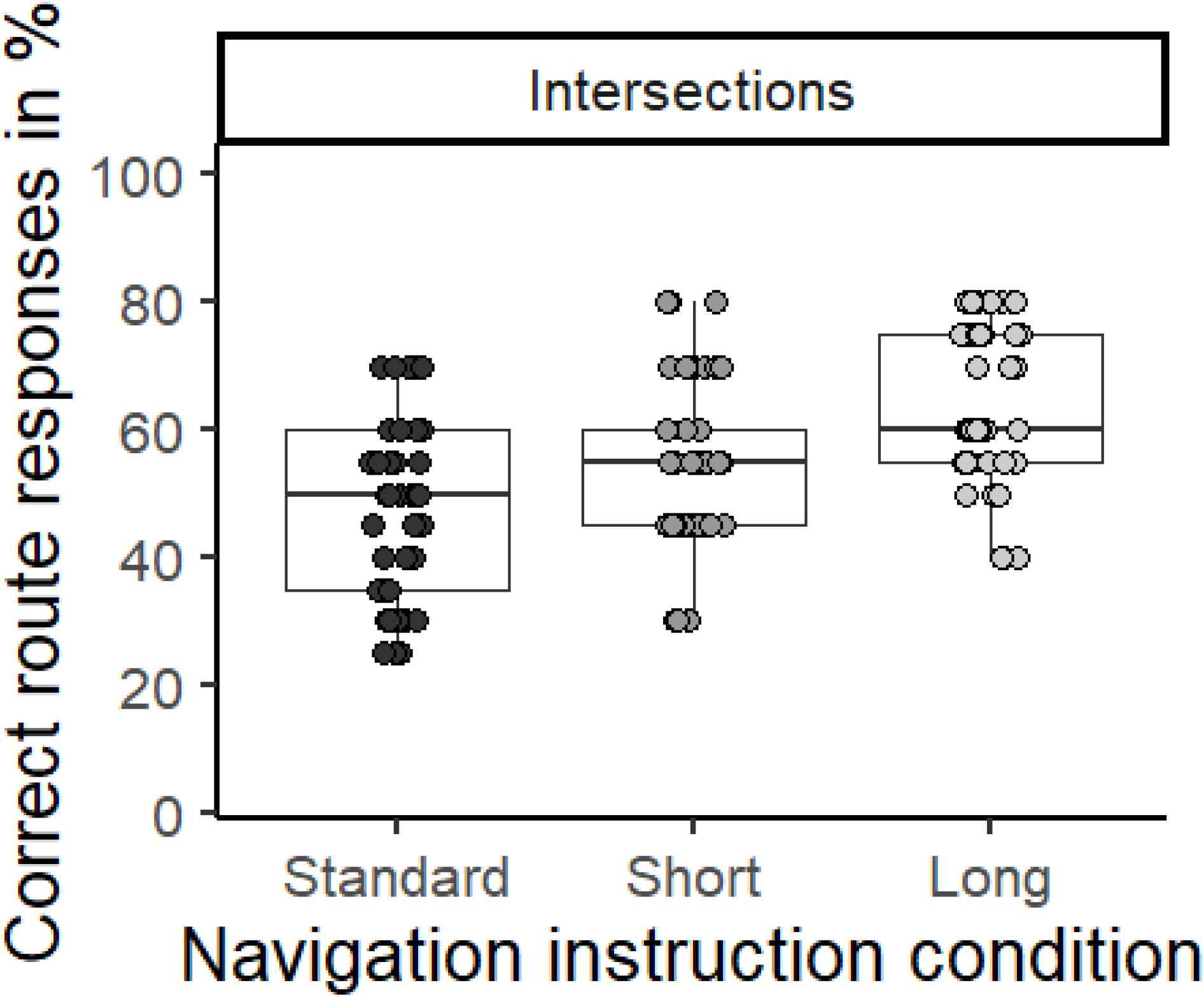
Boxplots and single subject performance scores of the correct route direction indication in the cued-recall task after video navigation. For correct route responses, 100% equals the correct response to the twenty landmarks at intersections. The performance measure could reach twenty different values with a step-width of 5%. Small dots represent individual values above the whiskers.

#### Route Knowledge

The ANOVA testing the percentage correct responses regarding landmarks at intersections revealed a significant effect of navigation instruction condition (*F*_(2,40)_ = 5.64, *p* = .007, *η*_p_^2^ = .220). Post-hoc comparisons revealed that the performance in the long navigation instruction condition was better (*M* = 64.7%, *SE* = 3.47%) than in the standard instruction condition (*M* = 48.2%, *SE* = 3.59%, *p* = .006). The short navigation instruction condition revealed intermediate performance (*M* = 53.9%, *SE* = 3.59%, *p*’s > .111). For individual route response performance, please see 3.

### Discussion

In Experiment 1, a paradigm shift from simulated driving to video-based pedestrian navigation was realized. This allowed us to test the effect of landmark-based navigation instructions on incidental spatial knowledge acquisition in a more realistic setting than simulated driving while allowing participants to remain seated as in the driving simulator study by Gramann et al. (2017). Two different landmark-based navigation instructions (short and long) were compared with a control group.

The incidentally acquired landmark knowledge was tested comparing the recognition sensitivity measure between the navigation instruction conditions. There was no significant main effect of the navigation instruction condition when the influence of the covariate perspective taking was controlled for. The interaction with the factor landmark type was also insignificant.

The missing replication of the positive impact of landmark-based navigation instructions on landmark knowledge during video navigation might be explained by the rather passive watching state of all participants. This seems to have diminished the attention allocation effects provoked by different auditory navigation instructions. As the video did not require any control of a primary task and walking speed was relatively slow, participants were able to fully concentrate on the instructions and the environment.

Possibly, differences in incidental spatial knowledge acquisition become only noticeable in actual dual-task conditions with the primary task (e.g., driving or real walking) requiring attentional resources. Furthermore, the task was interrupted at each intersection by presenting the figure with the three response keys. Stopping and masking the video content might have artificially suppressed the processing of the environment during intersection phases of the video navigation.

The results of incidentally acquired route knowledge revealed a positive impact of more detailed landmark-based navigation instructions. The long instructions condition outperformed the control group and tended to be better than the short version of the landmark-based instructions. This replicates the previous results of Gramann et al. (2017) and Wunderlich and Gramann (2018) even though new modifications of landmark-based navigation instructions were introduced. In addition, the navigation context had changed from simulated driving to an interactive video of pedestrian navigation that showed real-world visuals and included other traffic participants.

However, the new video-based setup was influenced by the individual perspective taking ability. The covariation with the PTSOT results might be due to the rather passive, video-based setup that allowed participants to process their surrounding environment with significantly more time as compared to simulated driving. At each turn, the video perspective changed whereas the physical reference frame of the navigator remained the same. Thus, individual differences in spatial abilities, like perspective taking or to some extend also the self-reported sense of direction, might have had a stronger influence on spatial knowledge acquisition during video watching. This needs to be considered carefully, when drawing conclusions about reliability and generalizability of the effects.

This first experiment replicated some of the results from previous studies but still lacked natural movement, the accompanying kinesthetic and proprioceptive information during real pedestrian navigation that might further support spatial knowledge acquisition of the surroundings. To allow for multimodal information integration of all senses involved in natural navigation through the real world, Experiment 2 implemented a real-world navigation task with pedestrians actively navigating the same route through Berlin.

## Experiment 2

In the second experiment, the experimental protocol changed from a stationary laboratory setup to a real-world setting allowing for free movement of navigators through their environment. This modification aimed at testing the ecological validity of the previously demonstrated incidental spatial learning effect during the use of long landmark-based navigation instructions in an unrestricted, real-world setting. The analysis of the brain activity accompanying assisted navigation was reported in a separate manuscript (Wunderlich & Gramann, 2020).

### Method

#### Participants

Of the initially acquired 35 data sets, the recordings of 22 participants (11 females) remained in the analysis after excluding all recordings with technical issues or of participants that had been familiar with 50% or more of the route. The remaining participants had a mean route familiarity of 9.52% (*SD* = 12.2%, standard: *M* = 11.5%, *SD* = 11.9%, long: *M* = 7.50%, *SD* = 12.8%, *F*_(1,20)_ = 0.59, *p* = .451). The age ranged between 20 and 39 years (*M* = 27.4 years, *SD* = 4.63 years). Each experimental condition consisted of eleven participants each with gender being balanced across conditions. Recruitment of participants was done using an existing database or personal contacts and reimbursement was received monetarily or by course credit. All participants had normal or corrected to normal vision and gave written informed consent prior to the study. The study was approved by the local ethics committee.

#### Procedure

Like the first experiment, this experiment took place in two sessions directly following each other. A short break of about 15min between the assisted navigation phase and the subsequent spatial tests was used for transporting the participant from the end of the route to the laboratory. The experiment lasted about 3 hours in total. During assisted navigation, participants navigated the identical route presented as video in Experiment 1. Two navigation instruction conditions were tested. The control group receiving standard navigation instructions and a second group receiving the long landmark-based navigation instructions from Experiment 1. Participants were unaware that they would be tested about the navigated environment after the navigation phase.

#### Navigation Phase

Each participant came to the Berlin Mobil Brain/Body Imaging Lab (BeMoBIL) to receive all necessary information about the experiment and to sign the informed written consent. They were then transported by car to the starting point of the route. After arrival, the EEG cap (eego, ANT Neuro, Enschede, Netherlands) was applied. Participants were instructed to follow the guidance of the auditory navigation instructions, be attentive to the surrounding traffic at all times, and stay where they were if they felt lost. An experimenter was following the participant during navigation to ensure that the participant would walk the correct route safely (see Figure 4a). The auditory navigation instructions were triggered manually by the experimenter at predefined trigger points alongside the route. For doing so, a customized browser-based application was developed which was controlled by a smartphone. Participant, as well as the experimenter, were provided with the auditory navigation instructions via Bluetooth headphones (Cheetah Sport In-Ear, Mpow, Hong Kong, China).

**Figure 4.**
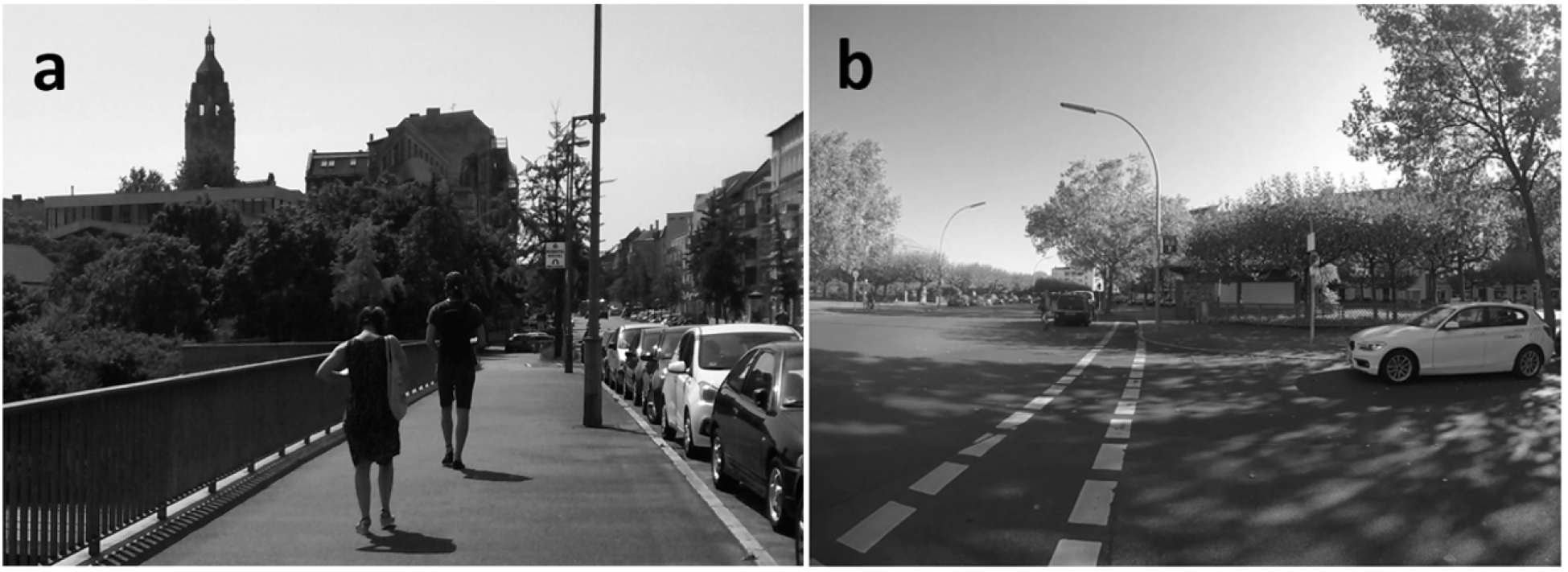
a) Real-world setup of the navigation task in Experiment 2. The experimenter followed the participant and initiated the auditory navigation instructions. b) Identical landmark pictures as in the cued-recall task of Experiment 1 were used in the cued-recall task of Experiment 2. This one shows a landmark at an intersection (“public restroom”) with a similar perspective as encountered during navigation (pictures were presented in color during the cued-recall task).

After navigation, participants rated their subjective navigation task-related load by answering the NASA-TLX (Hart, 2006; Hart & Staveland, 1988). Furthermore, participants responded to questions regarding route familiarity and in case they were part of the landmark-based instruction group, the acceptance of the modified navigation instructions.

#### Test Phase

In the subsequent spatial test session, participants were seated in a controlled laboratory setting at the BeMoBIL. A sketch map task as well as the cued-recall task had to be solved, followed by a digital version of the PTSOT (Hegarty and Waller, 2004, https://github.com/TimDomino/ptsot). During the tasks, participants were seated at a table and provided with paper and pen or computer screen and keyboard, respectively. Participants were still wearing the EEG cap in order to record brain activity during the spatial tasks as well.

The cued-recall task was identical with the one of Experiment 1, the same 60 screen captures from the video were used as landmark pictures including twenty pictures of each landmark type. Pictures of landmarks displayed their surroundings in the first-person perspective during walking (see Figure 4b). Every landmark picture was shown for three seconds and then a picture with the four possible response keys replaced the landmark picture until response. After each response, participants were asked to rate how confident they had been about their answer on a six-point scale ranging from ‘1 - very sure’ to ‘6 - I did not know’ or choose a seventh option stating ’I know that my last response was wrong’. Subsequent to the tasks, questionnaires were used to collect demographic characteristics, individual navigation preferences, and the self-reported SOD.

#### Statistical Analysis

A mixed measures ANOVA tested recognition sensitivity using the navigation instruction condition (standard, long) as between-subject factor and landmark type (intersections, straight segments) as within-subject factor. The percentage of correct responses for landmarks at intersections was tested by an one-factorial ANOVA with the between-subject factor navigation instruction condition (standard, long).

In case an individual measure correlated significantly with the dependent variable for each within-subject factor level, it was included as a covariate in the respective analysis. For additional information please see methods part of Experiment 1.

### Results

#### Individual Measures

Averaged across all participants, a mean error deviation angle in the PTSOT of 31.4° (*SD* = 24.5°, standard: *M* = 36.4°, *SD* = 30.2°, long: *M* = 26.5°, *SD* = 17.1°, *F*_(1,20)_ = 0.91, *p* = .352) and mean self-reported SOD of 3.73 (*SD* = 1.49, standard: *M* = 3.36, *SD* = 1.69, long: *M* = 4.09, *SD* = 1.22, *F*_(1,20)_ = 1.34, *p* = .261) was observed. Participants stated their subjective mental load after assisted navigation on a scale from 1 to 100 with *M* = 30.0 (*SD* = 18.5, standard: *M* = 22.6, *SD* = 16.2, long: *M* = 37.5, *SD* = 18.4, *F*_(1,20)_ = 4.04, *p* = .058).

The Spearman correlation of the angular error in the PTSOT showed a significant correlation with the percentage of correct responses to landmarks at intersections (*ρ* = −.504, *p* = .017, all other |*ρ*|’s < .297, *p*’s > .181). The correlation of the SOD rating and the percentage of correct responses to each landmark type was not significant (|*ρ*|’s < .337, *p*’s > .126). The subjective mental load correlated positively with the sensitivity d’ for landmarks at intersections (*ρ* = .457, *p* = .033, all other |*ρ*|’s < .334, *p*’s > .129). As the non-significant correlation of subjective mental load with the other factor level of recognition sensitivity, landmarks at straight segments, pointed in the opposite direction (*ρ* = −.270), subjective mental load did not fulfill the requirements for a covariate. Thus, only the PTSOT error was included as covariate in the one-factorial ANOVA testing the percentage of correct responses regarding landmarks at intersections.

#### Landmark Knowledge

Recognition sensitivity for landmarks at intersections and at straight segments was plotted in figure 5. When testing the recognition sensitivity, the main effect of landmark type (*F*_(1,20)_ = 4.22, *p* = .053, *η*_p_^2^ = .174) and the main effect of navigation instruction condition (*F*_(1,20)_ = 3.53, *p* = .075, *η*_p_^2^ = .150) did not reach significance. However, there was a significant interaction of both factors (*F*_(1,20)_ = 6.51, *p* = .019, *η*_p_^2^ = .246). The post-hoc comparisons of the interaction comparing the navigation instruction conditions for each landmark type revealed a significantly lower sensitivity for landmarks at intersections in the standard navigation instruction group (*M* = 1.47, *SE* = 0.25) compared to the landmark-based navigation instruction group (*M* = 2.47, *SE* = 0.25, *p* = .009).

**Figure 5.**
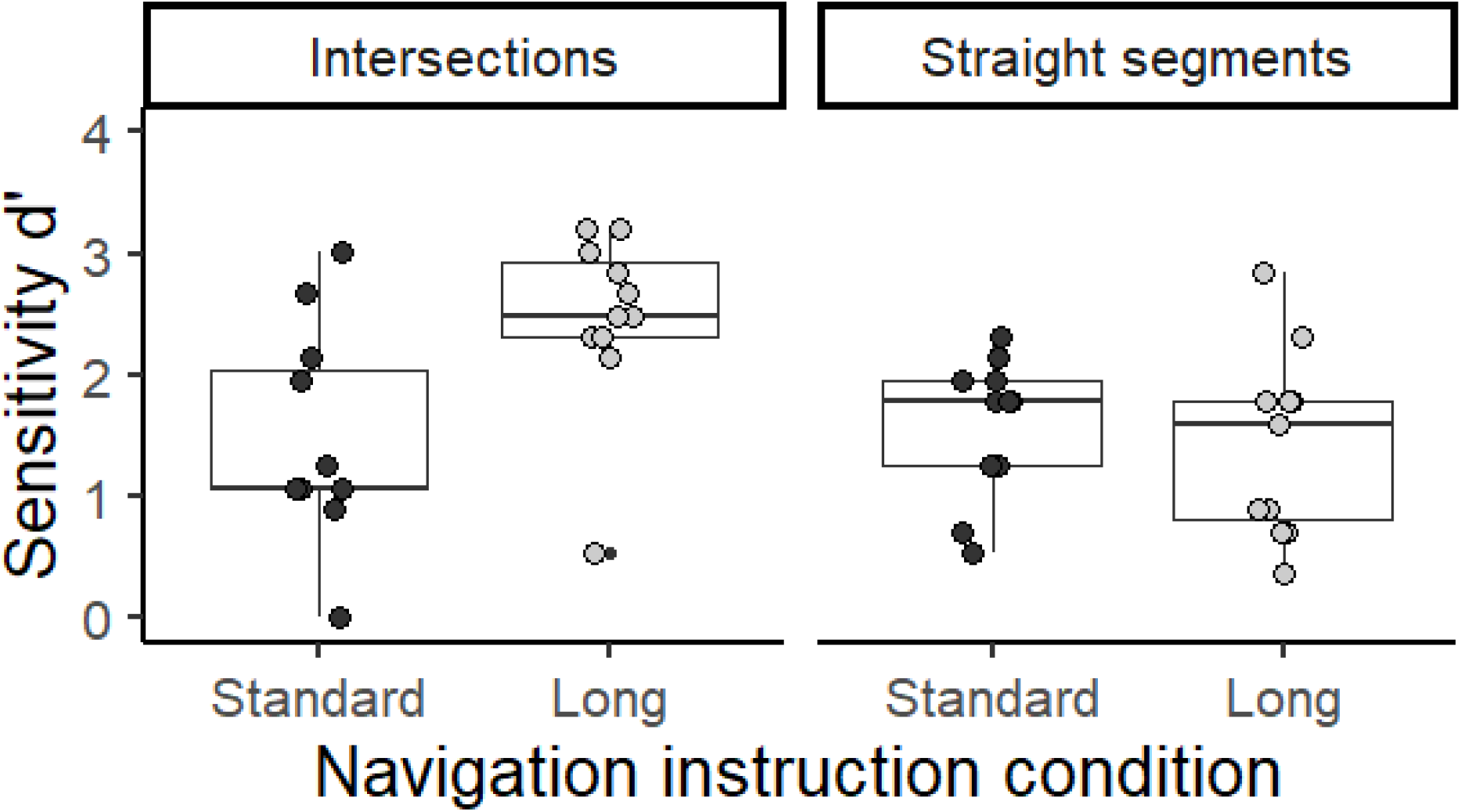
The sensitivity for correctly recognizing familiar landmarks was presented in box plots and single subject values of d’ for Experiment 2. The d’ represents the z-standardized rate of hits subtracted by the z-standardized rate of false alarms. Separated in columns, the sensitivity was plotted for landmarks at intersections and for landmarks at straight segments. Small dots represent individual values above the whiskers.

#### Route Knowledge

The ANCOVA for the percentage of correctly identified landmarks at intersections and associated navigation decision revealed a significant main effect of the covariate PTSOT error (*F*_(1,19)_ = 6.84, *p* = .017, *η*_p_^2^ = .265) as well as a significant main effect of the factor navigation instruction (*F*_(1,19)_ = 10.0, *p* = .005, *η*_p_^2^ = .346). Post-hoc comparisons revealed the control group to identify less landmarks correctly (*M* = 49.6, *SE* = 3.68) compared to the long landmark-based navigation instruction condition (*M* = 69.1, *SE* = 3.68). In figure 6, individual p erformance scores were presented.

**Figure 6.**
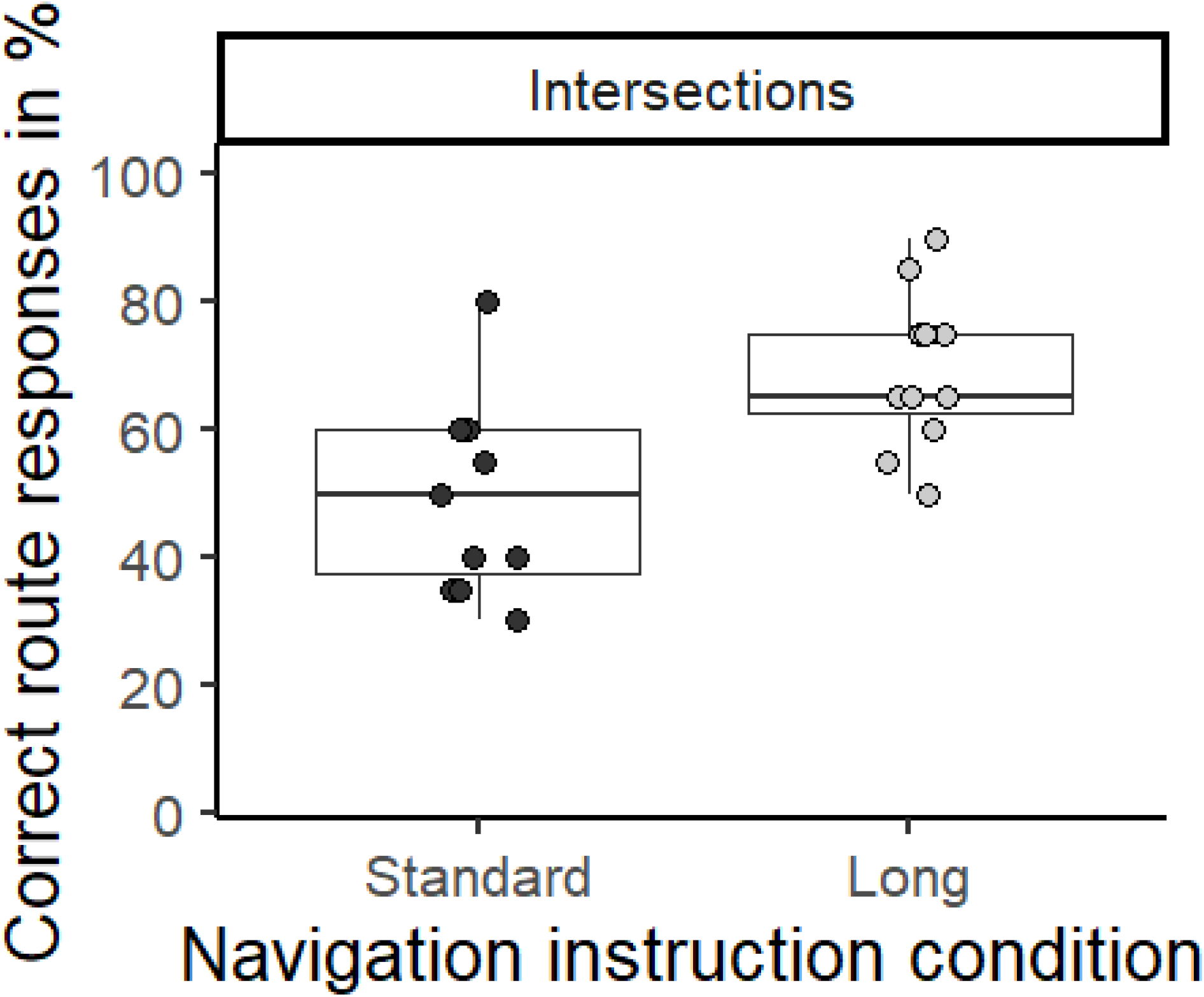
Boxplots and single subject performance scores of the correct route direction indication in the cued-recall task directly after assisted navigation through a real-world environment. For correct route responses, 100% equals the correct response to the twenty landmarks at intersections. Small dots represent individual values above the whiskers.

### Discussion

Experiment 2 tested the incidental spatial knowledge acquisition during assisted pedestrian navigation in an uncontrolled, dynamically changing real-world setting including cars and other pedestrians. Performance in the cued-recall task was tested as the second task after a short break following the assisted navigation phase. The results replicate previous results supporting the assumption that landmark-based navigation instructions enhance incidental spatial knowledge acquisition during assisted navigation.

In accordance with our hypothesis, the recognition sensitivity indicated enhanced spatial knowledge acquisition for landmark-based navigation instruction during navigation in the real world. The significant interaction effect of landmark type and navigation instruction conditions showed that participants receiving long landmark-based navigation instructions acquired more landmark knowledge at intersections than the control group.

When testing incidentally acquired route knowledge by considering only correct direction responses, the long instruction condition again outperformed the standard navigation instruction condition. Even though there was a significant correlation of correct responses for landmarks at intersections and the individual perspective taking ability (PTSOT), the covariate did not diminish the effect of navigation instruction conditions. Thus, it can be concluded that even though perspective taking ability seems to modulate spatial knowledge acquisition during navigation on a route level, an additional impact of landmark-based navigation instructions can be observed.

The hypothesized halo effect was not supported and a significantly higher recognition sensitivity for landmarks at intersections compared to straight segments was observed when using landmark-based navigation instructions. The landmark-based navigation instructions did not lead to a better recognition of other landmarks along the route in between intersections. This result contrasts with the results from experiment 1 which revealed a higher recognition sensitivity for landmarks at straight segments compared to landmarks at intersections for all tested instruction conditions. Whether this is due to the setup change from video-based real-world to unrestricted movement through the real-world was tested in the third experiment replicating the real-world pedestrian navigation scenario with an additional break between navigation phase and spatial tests.

## Experiment 3

The third experiment took place again in an unrestricted, real-life setting. The experimental setting was modified by extending the route and adding a two-week break between assisted navigation and testing the acquired spatial knowledge. This way, the third experiment aimed at replicating experiment 2 and testing the previously shown long-term effect of Wunderlich and Gramann (2018) in a setting providing full ecological validity. Additionally, the perspective of the landmark pictures in the cued-recall task was changed to a frontal view in order to test the quality of the incidentally acquired landmark and route knowledge.

### Method

#### Participants

In this experiment, 41 participants were recorded during real-world navigation. Due to technical issues or familiarity with 50% or more of the route, six of the participants had to be removed from the database. The remaining 35 participants (20 females) were aged between 20 and 34 years (*M* = 26.7 years, *SD* = 3.08 years) and stated their prior route familiarity with *M* = 15.4% (*SD* = 13.1%, standard: *M* = 17.9%, *SD* = 15.9%, short: *M* = 11.5%, *SD* = 9.03%, long: *M* = 17.0%, *SD* = 13.8%, *F*_(2,32)_ = 0.82, *p* = .448). Each participant was randomly assigned to an experimental condition (standard instructions: 8 female, 4 male; short instructions: 6 female, 6 male; long instructions: 6 female and 5 male participants). Participants were recruited through an existing database or personal contact and were reimbursed monetarily or received course credit. All had normal or corrected to normal vision and gave written informed consent prior to the study. The study was approved by the local ethics committee.

#### Procedure

The experiment was scheduled in two sessions separated by a break of two weeks. Taken together, both experimental sessions lasted about 3 hours and 30 minutes. In the first part, participants navigated along a predefined route through an urban environment, the district of Charlottenburg, Berlin in Germany. The route overlapped with the route in Experiment 1 and 2. Each participant was guided by auditory navigation instructions. There were three groups of participants including a control group who received standard navigation instructions. The landmark-based navigation instruction groups were either receiving the short, or long landmark-based navigation instructions. After a break of two-weeks (13 to 17 days, *M* = 14.1 days, *SD* = 0.73 days), the participants were invited to return and to solve spatial tasks in a laboratory setting. Participants were informed about the second session without providing information about its content. The relatively long break between navigation and test phase allowed for investigating the generalizability of the previously reported long-term impact of landmark based navigation instructions on spatial knowledge acquisition (Wunderlich & Gramann, 2018). Importantly, and in contrast to Wunderlich and Gramann (2018), participants walked through a city environment during the navigation phase and were also moving through the same city during their daily activities between navigation and test phase. Thus, they were probably experiencing a similar city environment that interfered with spatial knowledge about the experimental route. This way, the experiment represents an ecologically valid case of long-term spatial knowledge acquisition during assisted navigation.

#### Navigation Phase

An experimenter met each participant at a train station and guided them to the starting point of the route. There, participants were informed about the upcoming navigation task. They were instructed to follow the auditory navigation instructions, to pay attention to the traffic while crossing streets, and to stop in case they did not know which direction to proceed. While navigating the route, an experimenter was walking in the vicinity of the participant to intervene in case of hazards and ensure that the participant would follow the correct route (see Figure 4a). The experimenter also manually triggered the auditory navigation instructions at predefined trigger points alongside the route using a browser-based application on a smartphone. The auditory navigation instructions were provided to both, the participant and the experimenter, via Bluetooth headphones (Cheetah Sport In-Ear, Mpow, Hong Kong, China). The route consisted of 40 intersections including 22 direction changes. Participants arrived at the end of the route after walking approximately 60 minutes.

Following the navigation phase, navigators provided their subjective rating of the navigation task-related load as assessed using the NASA-TLX (Hart, 2006; Hart & Staveland, 1988). In addition, participants answered questions about route familiarity and acceptance of the modified navigation instructions.

#### Test Phase

During the spatial test session in a controlled laboratory setting, the cued-recall task and a video-turn task were recorded. Before the two tasks, participants were equipped with EEG (BrainAmps, Brain Products, Gilching, Germany) and seated at a table in front of a computer screen and keyboard. Analog to the cued-recall task of Experiment 1 and 2, 120 landmark pictures were presented including forty pictures of each landmark type. Pictures displayed their surroundings but in contrast to experiments 1 and 2, landmarks were presented in a front view perspective (see Figure 7b compared to Figure 1b and Figure 4b). Novel landmark pictures were photographs taken in a similar way and similar weather conditions in a neighboring area. Every landmark picture was shown until the participant responded by pressing one of the four possible arrow keys. After each response, participants were asked to rate how confident they had been about their answer on a six-point scale ranging from ‘1 - very sure’ to ‘6 - I did not know’. After the tasks, questionnaire data was collected including demographic characteristics, individual navigation styles, and self-reported SOD were assessed.

**Figure 7.**
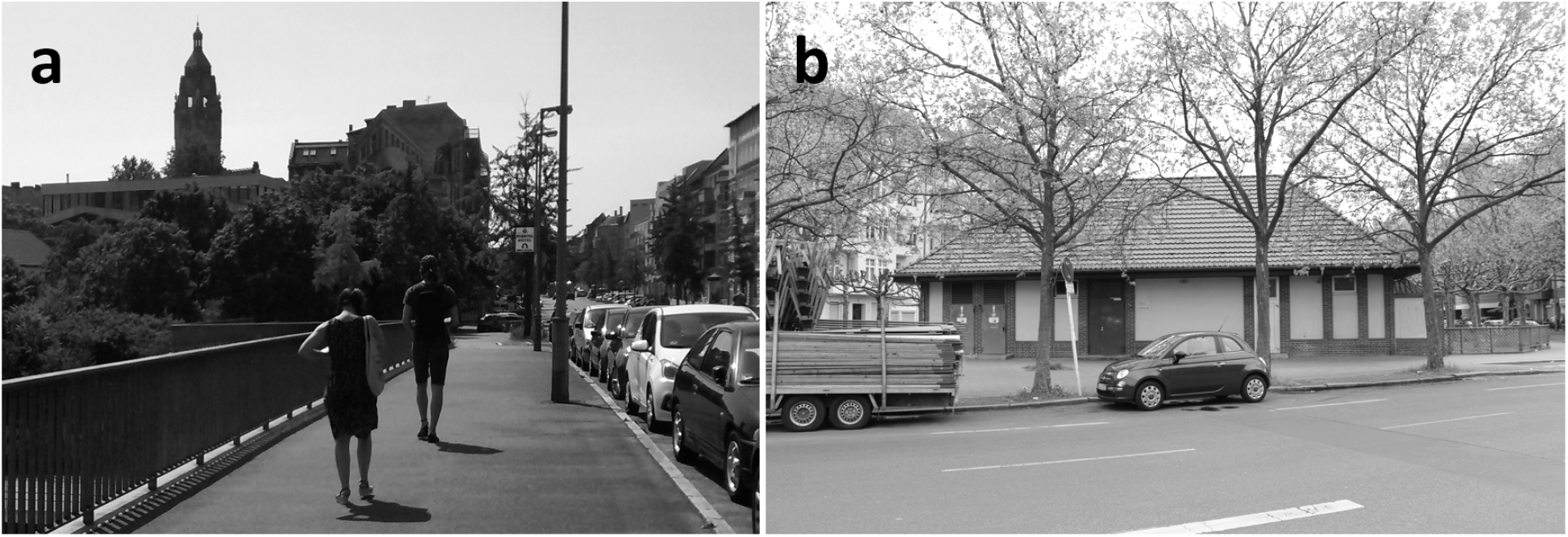
a) Real-world setup of the navigation task in Experiment 3. The experimenter followed the participant and initiated the auditory navigation instructions. b) Frontal view picture of a navigation relevant landmark (“public restroom”) as used in the cued-recall task (pictures were presented in color).

#### Statistical Analysis

As in Experiment 1, two analyses of variance were computed to test whether landmark-based navigation instruction lead to better landmark and route knowledge. Investigating the acquired spatial knowledge, a mixed measures ANOVA tested the recognition sensitivity with the between-subject factor navigation instruction condition (standard, short, long) and the repeated measures factor landmark type (intersections, straight segments). Additionally, the percentage of correct responses for landmarks at intersections was tested in a one-way ANOVA comparing the three navigation instruction conditions. For additional information, please see methods part of Experiment 1.

### Results

#### Individual Measures

Participants rated their sense of direction with *M* = 4.29 (*SD* = 1.74, standard: *M* = 3.33, *SD* = 1.67, short: *M* = 5.17, *SD* = 1.64, long: *M* = 4.36, *SD* = 1.50, *F*_(2,32)_ = 3.91, *p* = .030) with values of short being significantly higher than standard (*p* = .026, other *p*’s > .404).

The subjective mental load during assisted navigation was on average 15.0

(*SD* = 9.32, data missing for 14 participants). The Spearman correlations of the SOD rating and the subjective mental load with the dependent variables were not significant (|*ρ*|’s < .239, *p*’s > .300). Thus, neither the SOD rating nor the subjective mental load were included as covariate in the analyses.

#### Landmark Knowledge

Individual and group values of recognition sensitivity are presented in figure 8. The ANOVA testing the recognition sensitivity revealed a main effect of landmark type (*F*_(1,32)_ = 142, *p* < .001, *η*_p_^2^ = .817) and a main effect of navigation instruction condition (*F*_(2,32)_ = 5.31, *p* = .010, *η*_p_^2^ = .249). In addition, the interaction of both factors was significant (*F*_(2,32)_ = 4.07, *p* = .027, *η*_p_^2^ = .203). The recognition sensitivity for landmarks at intersections (*M* = 1.66, *SE* = 0.12) was significantly higher compared to landmarks at straight segments (*M* = 0.20, *SE* = 0.14, *p* < .001). The control group had significantly lower d’ (*M* = 0.33, *SE* = 0.19) than the landmark-based navigation instruction conditions (short: *M* = 1.13, *SE* = 0.19, *p* = .017; long: *M* = 1.06, *SE* = 0.20, *p* = .039). Short and long navigation instruction conditions were comparable (*p* > .999). The post-hoc comparisons of the interaction comparing the navigation instruction conditions for each landmark type revealed a lower recognition sensitivity for landmarks at intersections in the standard navigation instruction group (*M* = 0.90, *SE* = 0.21) compared to the landmark-based navigation instruction groups (short: *M* = 1.98, *SE* = 0.21, *p* < .001; long: *M* = 2.10, *SE* = 0.22, *p* < .001). All other *p*’s were above .093.

**Figure 8.**
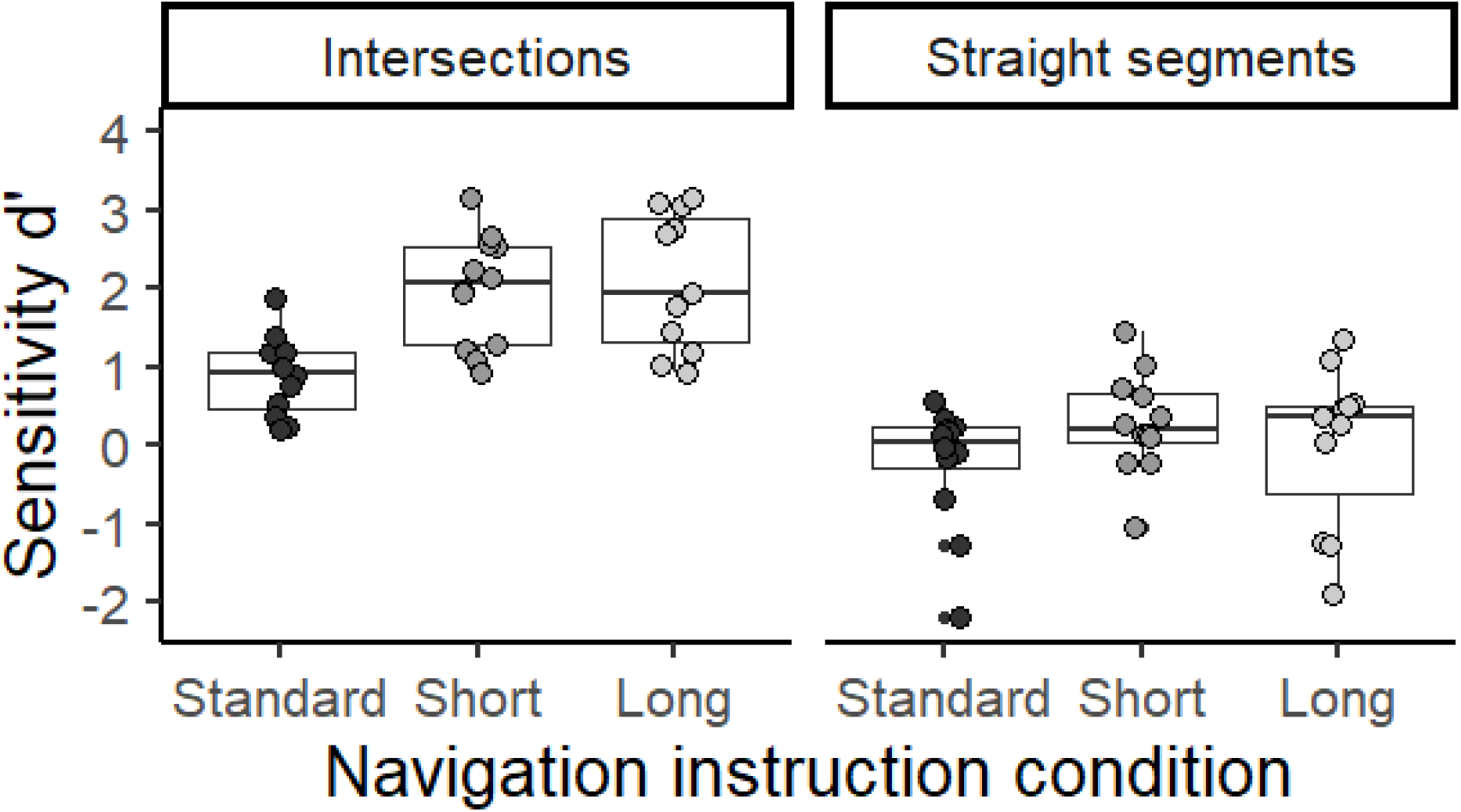
The sensitivity for correctly recognizing familiar landmarks was presented in box plots and single subject values of d’ for Experiment 3. The d’ represents the z-standardized rate of hits subtracted by the z-standardized rate of false alarms. Separated in columns, the sensitivity was plotted for landmarks at intersections and for landmarks at straight segments. Small dots represent individual values above the whiskers.

#### Route Knowledge

The one-way ANOVA comparing the percentage of correct responses to the landmarks at intersections between the three navigation instruction conditions revealed a significant difference (*F*_(2,32)_ = 6.07, *p* = .006, *η*_p_^2^ = .275). The standard navigation instruction condition had a significantly lower performance (*M* = −30.2, *SE* = 3.45) as compared to the landmark-based navigation instruction condition (short: *M* = 42.9, *SE* = 3.45, *p* = .041; long: *M* = 46.6, *SE* = 3.60, *p* = .007). Performance of short and long navigation instruction condition demonstrated comparable results (*p* > .999). For individual performance scores, please see figure 9.

**Figure 9.**
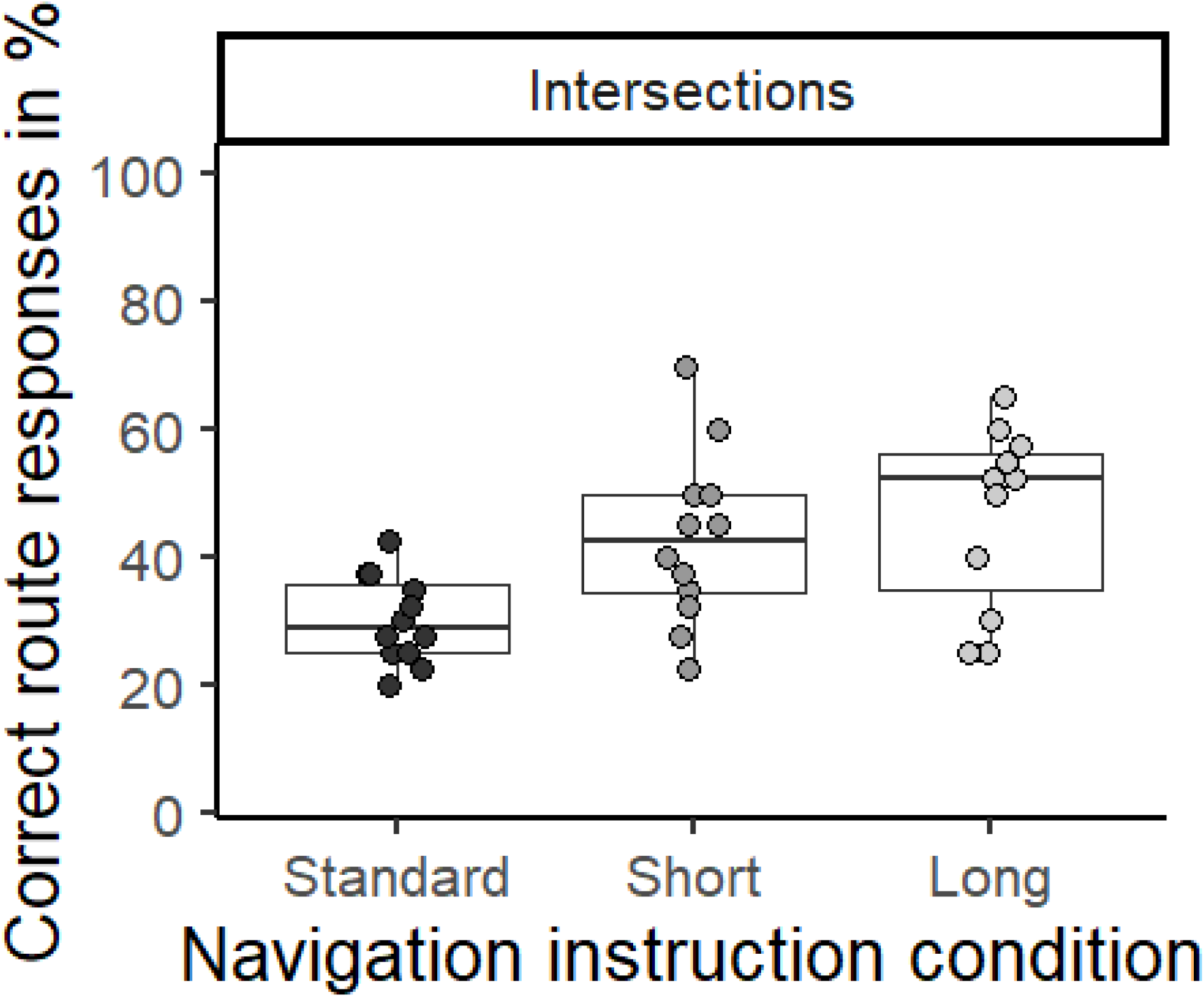
Boxplots and single subject performance scores of the correct route direction indication in the cued-recall task directly two weeks after assisted navigation through a real-world environment. For correct route responses, 100% equals the correct response to the forty landmarks at intersections. Small dots represent individual values above the whiskers.

### Discussion

Experiment 3 followed-up on Experiment 2 investigating incidental spatial knowledge acquisition during assisted pedestrian navigation in the real-world. Even after a break of two weeks, the results of the cued-recall task demonstrated an improved recognition sensitivity when navigating with landmark-based navigation instructions.

When investigating the recognition sensitivity with signal-detection theory, we observed improved performance for landmarks at intersections for landmark-based as compared to standard navigation instructions. Landmark-based navigation instructions also led to better route knowledge as reflected in the percentage of correct direction responses. However, long and short landmark-based instruction did not differ with regards to route knowledge. It can thus be concluded that after a break of two-weeks the advantage of long versus short navigation instructions as seen in the video-based results disappears and both navigation instructions perform better than standard instruction but comparably regarding the successful recollection of landmark and route knowledge.

As in Experiment 2, no halo effect of the landmark-based navigation instructions were observed as the recognition sensitivity was significantly higher for the augmented landmarks at intersections compared to salient objects at straight segments. In contrast to all previous experiments, landmark pictures were presented from a frontal view instead of a navigators’ perspective when navigating through the environment. This might have been especially detrimental to the recognition of landmarks positioned at straight segments as participants might not have turned their heads to inspect all buildings alongside the route and thus, landmarks at straight segments were less likely encountered from this perspective. When approaching an intersection or crossing a street turning the head to left and right is a well-trained behavior essential for maintaining safety. That is why, participants of all navigation instruction conditions likely have processed landmarks at intersections from more than one perspective leading to better recognition performances.

A limitation of experiment 3 is missing data regarding the individual measures. The subjective task load was only collected from a subset of the sample and the individual perspective taking ability was not tested. In experiment 1 and 2, a significant impact of individual perspective taking on acquired landmark or route knowledge was shown. However, this impact did not explain or diminish the effect of navigation instruction conditions.

To sum up, when pedestrians use landmark-based navigation instructions in the real-world environment, they do acquire more spatial knowledge about the travelled environment especially about the auditory augmented landmarks at intersections. This advantage is still measurable after two weeks replicating the long-term effects previously shown in simulated driving through a sparse and controlled virtual world by Wunderlich and Gramann (2018). This long-term improvement takes place even when participants commute through a similar city environment in the time period between navigation and test phase potentially interfering with memory consolidation.

## General Discussion

Three experiments investigated incidental spatial knowledge acquisition during assisted pedestrian navigation in ecologically valid scenarios. It was investigated whether there is evidence for enhanced incidental spatial knowledge acquisition when using different versions of landmark-based navigation instructions and whether these also change the processing of the environment in general. To that end, three experiments were conducted investigating the performance in a cued-recall task as an indicator for landmark- and route-level spatial knowledge acquired during a single exposure to an unfamiliar environment. The navigation contexts used were videos of (Experiment 1) as well as real-world pedestrian navigation (Experiment 2 and 3) through an unfamiliar city area. In summary, enhanced incidental spatial knowledge acquisition was shown for landmark-based navigation instructions in all three experiments. But there was no hint of a generalization of spatial knowledge acquisition beyond the augmented landmarks to other salient objects in the environment. Thus, no effect of landmark-based navigation instructions reflecting a generally increased processing of the environment was observed. Individual measures were controlled for and did not impact the effects of the navigation instruction conditions on spatial knowledge acquisition.

Following the argument that decreased spatial memory performance associated with the use of navigation aids is likely caused by divided attention between the control of movement and the navigation assistant (Gardony et al., 2013; Gardony et al., 2015), spatial knowledge acquisition was expected to be impaired when using standard navigation instructions irrespective of the landmark type. In the studies of Gardony and colleagues, only the retrieval of landmarks at intersections was investigated. Here, comparing spatial knowledge for different landmark types, we still found poorer landmark knowledge at intersections for the standard navigation instruction condition while having comparable landmark knowledge for landmarks at straight segments.

The low spatial knowledge acquisition was likely caused by instructions biasing the attention towards the intersection while landmark-based navigation instructions helped to process environmental features. The augmentation of landmarks allowed to encode a specific landmark in association with a turning decision.

The results of all three experiments replicated the positive impact of landmark-based navigation instructions on spatial knowledge acquisition which was previously shown for visual landmark augmentation (Krukar et al., 2020; Li et al., 2014; Löwen et al., 2019; Tom & Denis, 2003). Furthermore, the reported experiments replicated findings from Gramann et al. (2017) and Wunderlich and Gramann (2018) demonstrating an increased cued-recall performance especially regarding landmarks at intersections after navigating with auditory landmark-based navigation instructions. This underpins the robustness and the ecological validity of the reported incidental spatial learning effect when using landmark information in auditory navigation instructions.

The acquired landmark and route knowledge did not depend on individual spatial abilities demonstrating that landmark-based navigation instructions improve spatial knowledge acquisition for all navigators irrespective of their spatial abilities. Even though an impact of individual perspective taking ability on spatial knowledge acquisition was revealed, this effect did not explain the increased spatial knowledge acquisition based on the landmark-based navigation instructions.

In addition, the results provided evidence that adding detailed information about a landmark in addition to a landmark name is similar to the personal preference navigation instruction condition used in Gramann et al. (2017) and Wunderlich and Gramann (2018) regarding spatial knowledge acquisition. In Experiment 1, only an advantage of long landmark-based navigation instructions was found for incidental route knowledge acquisition. It was beneficial to add semantic information in the auditory navigation instructions. Only naming the landmark was not sufficient. It can be argued that the detailed information about the landmarks in long navigation instructions may have helped to identify the landmark and this way eased the landmark recognition in the cued-recall task. However, the additional semantic information was mostly related to the function of the object and did not include characteristics that were visible (e.g., “Go straight at the Malteser building. This aid organization counts one million members.”). As such, the identification of landmarks was similar in both landmark-based navigation instructions and a confounding effect can be ruled out. In Experiment 3 testing spatial knowledge after a break of two weeks, no difference between the landmark-based navigation instructions was measurable, but both modifications significantly improved the incidentally acquired spatial knowledge compared to the control group. This replicates the driving simulator studies (Gramann et al., 2017; Wunderlich & Gramann, 2018) and hints at a significant impact of landmark-based instruction but only a rather short-term advantage of the additional landmark information when tested in real-world pedestrian navigation.

Another hypothesis based on the brain activity results of Wunderlich and Gramann (2018) was that landmark-based instructions lead to a generally increased allocation of attention towards the environment. This would be supported when the navigation instructions at relevant navigation points like intersections also impacted how the surroundings were processed during straight segments where no navigation instructions were provided. The comparable recognition rates for landmarks at straight segments across navigation instruction conditions in all experiments, however, did not support such a halo-effect of landmark augmentation.

An interpretation of all reported results is still restricted to the investigation of acquired spatial knowledge when navigating one route in an unfamiliar environment and thus primarily to the acquisition of landmark and route knowledge. A generalization to spatial knowledge acquisition based on multiple uses of landmark-based navigation instructions for the same route or different routes with overlap is not yet possible. However, the results let us expect that spatial knowledge acquisition would be enhanced even more. Multiple use of landmark-based navigation assistance in the same area would also ease the assessment of acquired survey knowledge. It was shown that landmark, route and survey knowledge develop in parallel, rather than in a sequential fashion revealing increasing convergence with increasing exposure to the same environment (Buchner & Jansen-Osmann, 2008; Kim & Bock, 2020). Fostering the acquisition of survey knowledge would be the ultimate goal of a learning-oriented navigation assistance systems as survey knowledge would allow for more flexible use of the acquired spatial knowledge (e.g., short cuts). For now, the cued-recall performance allows only for conclusions about the acquired landmark and route level knowledge, but not survey knowledge. Whether the landmark-based navigation instructions can help to improve survey knowledge after repetitive use of the system in the same environment has yet to be tested.

In summary, this series of experiments demonstrates improved spatial knowledge acquisition including both the transfer from low to high realism of the environment as well as the realism of the movement through the environment and allowing for interfering navigation experiences. Despite all setup changes, the results indicate that landmark-based navigation instructions lead to improved spatial knowledge acquisition compared to standard navigation instructions that are currently used as default in available navigation assistance systems.

## Conclusions

The findings of the three experiments reported here replicated the previously described increased incidental spatial knowledge acquisition associated with landmark-based navigation instructions. This effect was shown to be generalizable across different navigation contexts and types of locomotion. Thus, landmark-based navigation instructions are a promising tool to keep the processing of the environment high and foster the extraction of navigation relevant information. Future research should address the multiple use of landmark-based navigation instructions and the accompanying spatial knowledge acquisition. This way, landmark-based navigation assistants may train the users’ orientation abilities enabling them to autonomously navigate the environment in the future.

## Acknowledgments

This research was supported by a stipend from the Stiftung der Deutschen Wirtschaft to AW. We give thanks to André Brandewiede, Maike Fischer, Anja Marckwardt (sorted by lastname) as well as the students in the neuroergonomics project courses in the year 2017 and 2018 for helping to prepare and conduct the experiments.

## Data Availability

Data were collected at TU Berlin. The data of single or all experiments are available on request from the corresponding author AW.

## Key Points

- Landmark-based navigation instructions enhance the incidental acquisition of landmark and route knowledge during assisted navigation.
- This improved incidental spatial knowledge acquisition was tested in different experimental setups and was shown to be robust and ecological valid.

## Context

Gramann started the research series investigating incidental spatial knowledge acquisition during assisted navigation in 2014 using simulated driving. Wunderlich followed up on this first experiment (Gramann et al., 2017) in her Master’s Thesis (Wunderlich & Gramann, 2018) which then was further developed as the quest for her PhD under constant supervision of Gramann. Gramann had coined the term natural cognition and with establishing the Berlin Mobile Brain/Body Imaging Lab, he built an research environment where the brain activity of freely moving and interacting participants can be investigated with the least amount of restrictions possible. Within the navigation project the authors even left the lab to reach out for the neurophysiological mechanisms underpinning spatial cognition in general and incidental spatial knowledge acquisition in particular using the highest ecological valid setup.

